# Temporal coding enables hyperacuity in event based vision

**DOI:** 10.1101/2025.03.25.645190

**Authors:** Eldad Assa, Alexander Rivkind, Michael Kreiserman, Fahad Shahbaz Khan, Salman Khan, Ehud Ahissar

**Author notes:** Equal contribution (Eldad Assa, Alexander Rivkind).

## Abstract

The fact that the eyes are constantly in motion, even during ‘fixation’, entails that the spike times of retinal outputs carry information about the visual scene even when the scene is static. Moreover, this motion implies that fine details of the visual scene could not be decoded from pure spatial retinal representations due to smearing. Understanding the interplay of temporal and spatial information in visual processing is thus pivotal for both biological research and bio-inspired computer-vision applications. In this study, we consider data from a popular event-based camera that was designed to emulate the function of a biological retina in hardware. Similarly to biological eye, and in contrast to standard frame-based cameras, this camera outputs an asynchronous sequence of “spike” events. We used this camera to obtain dataset of event streams of tiny images, i.e., images whose recognition is impaired by photosensor’s pixelization and thus their recognition requires hyperacuity. Using these datasets we demonstrate here the superiority of event-based spatio-temporal coding over frame-based spatial coding in the recognition of tiny images by artificial neural networks (ANNs). We further demonstrate the benefits of event sequences for unsupervised learning. Interestingly, Vernier hyperacuity, which is a standard measure of shape hyperacuity, emerged in ANNs following training on tiny images, resembling the natural hyperacuity observed in humans. Our findings underscore the essential role of precise temporal information in visual processing, offering insights for advancing both biological understanding and bio-inspired engineering of visual perception.

## 1 Introduction

Human vision is known for its remarkable spatial resolution, often exemplified by visual hyperacuity—a capability that enables humans to perceive fine details in small or distant objects with precision exceeding the spatial resolution of retinal photoreceptors [1, 2]. This skill is evident in tasks like vernier’s, where individuals can detect minuscule misalignments between two lines, even when the offset is measured in a few arcseconds—a scale significantly smaller than the angular separation corresponding to the smallest photoreceptors [3]. The underlying mechanisms that allow such extraordinary precision, given the relatively coarse structure of the retina, are not yet understood.

Visual acuity is known to improve with longer fixations, a phenomenon documented across several decades of research [4, 5, 6, 7]. During fixation, the eyes drift within a region of interest in various directions at instantaneous speeds ranging around 1 to 10 deg/s and amplitudes of several arcminutes [8]. Retinal cells respond to changes in illumination caused by the interaction between the eye’s motion and local image contrasts during this fixational drift [9, 8]. This active sensing leads to a rapid loss of spatial correspondence between image details and retinal activations [10], a phenomenon known as visual smearing [11]. Uncovering the mechanisms that allow the visual system to generate clear perception and overcome smearing caused by either external motion or eye motion is crucial to the understanding of dynamic visual perception.

It has been hypothesized that the visual system may counteract this smearing through spatial interpolation [1, 12] or statistical inference [13], or bypass it by processing temporal delays between retinal spikes [9, 14, 15, 16] as the temporal component of spatio-temporal filters [17]. In particular, the usage of precise spike timing to encode information in biological neuronal networks gained significant empirical and theoretical support over the last decades [18, 19]. Specifically, it was shown that precise spike timing can be used to extract spatial information from static visual stimuli [8, 14], and in the context of vernier offsets it was shown that a temporal difference could be perceived as a spatial offset [20, 1, 17], while best performance in humans was demonstrated with combined spatio-temporal cues [21]. These plausible mechanisms for maintaining high acuity by the visual system despite continuous retinal motion, may also provide insights into how artificial vision systems might improve their acuity beyond the resolution of the cameras’ photosensor array.

In a parallel field of research of biologically inspired computer vision, an event-based (EB) visual sensor was designed according to the architecture of the biological retina in order to emulate its function [22]. The motivation behind the development of this sensor, also known as ‘silicon retina’, was both to use it as a model for visual processing in the brain, and to gain an electronic sensor that encompasses the advantages of the biological retina over standard camera sensors (e.g. high dynamic range, low energy consumption, computation with low precision elements) [22, 23]. Current standard vision systems are typically based on frame-based (FB) cameras, which differ from biological systems as they collect information synchronously across all photosensors at a fixed rate; this leads to inefficient representation as redundant data are transmitted for regions of the environment where changes are slow, while information in regions where changes are faster than the acquisition rate is missing or distorted. In contrast, and similarly to the biological retina, EB cameras generate asynchronous events only when a photosensor experiences a change in luminance [24]. Each such event contains the precise timestamp of the event, the identity of the photosensor at which the event occurred (its x,y position on the sensor), and the polarity of the event (whether the luminance increased or decreased). EB sensors were used to model biological visual processing [25, 26], and their straightforward advantages were demonstrated in various applications, including visual recognition [27, 28, 29, 30, 31], image reconstruction [32, 33, 34, 35], 3D reconstruction [36, 37, 38, 39], visual tracking [40, 41, 42, 43], SLAM [44, 45, 46] and robotics and control [47, 48, 49, 50]. In contrast, although it was shown that the precise event’s timestamp contains information about the spatial pattern of the image [51, 52], the potential advantage of precise temporal coding for image recognition was rarely explored [29, 53, 31]. This is mainly due to two factors: 1) in most EB image recognition models events are first binned in time, resulting in frame-like representation which already loses precise time information [54]; 2) popular EB datasets for image recognition, such as the pioneering [55], contain sufficient spatial information, which precluded revealing the advantage of precise timing [53]. For a comprehensive survey of EB vision see [54] and [56].

The current work aims at bridging this gap. Specifically we show how utilizing the temporal information embedded within the visual EB data facilitates recognition of tiny still images, such as digits or fashion items. We opted for tiny, barely recognizable, images to specifically explore how precise temporal information, acquired during active vision [9], can be used to achieve hyperacuity.

Our experiments show that even images that are seemingly unrecognizable due to pixelization by the photosensor array, are classified surprisingly well using accurate temporal information and an appropriate classification model. We demonstrate this by limiting the availability of temporal data, which results in performance degradation. We also collected traditional, frame-based (FB) datasets for control, and demonstrated the advantage of EB sensing in classification accuracy in most of the settings. Furthermore, the essential relative motion and continuous acquisition of EB sensing allowed exploiting the constancy of the stimulus during each perceptual epoch to augment training, and train in an unsupervised and contrastive manner, demonstrating additional advantage for EB coding.

Investigating our model’s internal representations further, we show that training for image recognition in our demanding, near threshold, setting leads to the emergence of Vernier hyperacuity akin to that of humans. Our results highlight the significance of temporal coding as both a fundamental aspect of biological vision and a potentially beneficial component in computer vision. Additionally, we contribute a novel bio-inspired visual dataset, offering a benchmark where, for the first time, the fine temporal dynamics have a profound effect on image recognition performance.

## 2 Results

### 2.1 Creating tiny event-based versions of popular datasets

We used an EB camera (DAVIS240C, iniVation AG; [57]) to simulate the retina’s response to small static objects sampled by sensor motion. The setup was used to convert 3 popular FB datasets (MNIST [58], Fashion-MNIST [59] and Kuzushiji-MNIST [60]) to EB datasets of tiny symbols. Equivalent FB datasets were acquired for each condition (source dataset x distance) in a separate acquisition session, using the same camera in FB mode (the DAVIS240C sensor produces both events and frames). The FB tiny-symbol datasets were collected with the camera being still and the image falling on the center of the silicone retina.

In total, six EB datasets and six FB datasets were generated, corresponding to three source datasets and two distances. Their names are composed of the short name of the source dataset (MNIST, FMNIST and KMNIST) with ‘ebt’ (EB tiny) or ‘fbt’ (FB tiny) prefix and ‘-D1’ or ‘-D2’ suffix to signify camera’s distance (for example, event based dataset based on MNIST and generated with distance D2 is denoted by ebtMNIST-D2 and so on). The ratio between the pixel-size on the sensor and on the display was estimated to be between 1:7 to 1:8 for D1 and between 1:12 to 1:13 for D2 (see Methods).

Each such dataset includes all the 60K training samples and 10K test samples from the source datasets. The setup and few examples of events and frames acquired in different conditions are depicted in Figure 1A,D,E,F. As can be seen in the Figure, the tiny symbols are hardly recognizable by the human eye when examining both the event streams and the individual frames (Fig.1E,F,I).

**Figure 1:**
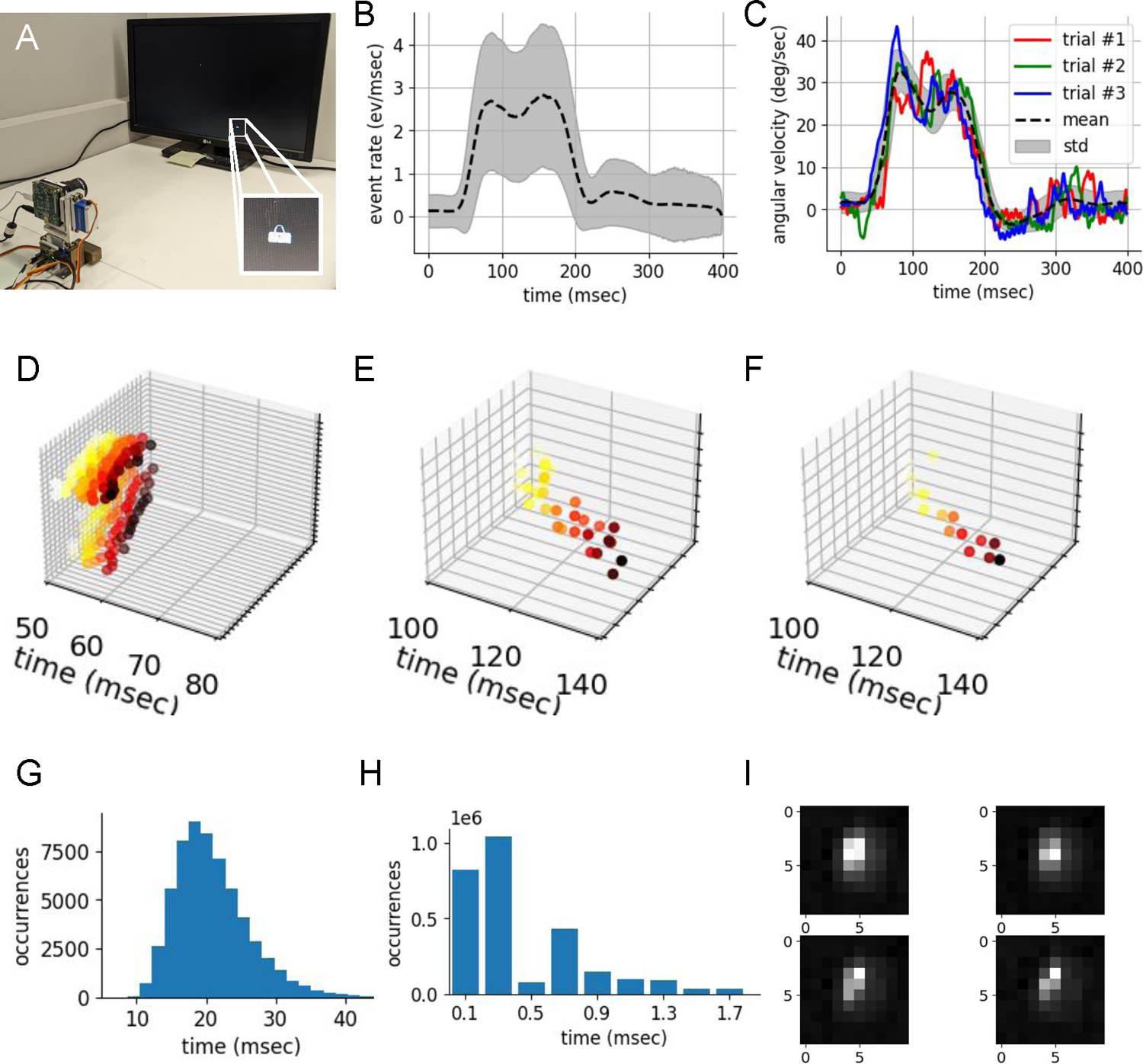
Experimental setup and data collected. **A A photo of the setup that was used to acquire the tiny images datasets**. The setup is composed of an EB camera mounted on a small robotic device which allows it to rotate, and a computer screen on which the symbols are displayed. Symbols are displayed at their original resolution of 28×28 pixels as seen in the inset on the right-bottom corner. **B Event rate (events/ms) during single horizontal rotation**. ebtMNIST-D1 dataset acquisition (MNIST samples at distance D1). Mean ± STD over 60K training-set samples. **C Horizontal angular velocity of the camera during ebtMNIST-D1 dataset acquisition**. Black dashed line and gray area correspond to mean and STD over 60K training samples respectively; color traces (red, blue and green) correspond to traces of 3 single-sample acquisitions. B-C) Time ‘0’ corrsponds to motor command initiation. **D,E,F) 3D (x,y,t) raster plots of events acquired during a single motion while MNIST symbol was displayed in front of the camera**. Time axis is marked while the orthogonal 2d plane represents the 2d sensor’s pixels matrix plane; points’ color corresponds to events’ timestamp (‘t’ value), events occurring later are darker. Only positive events are plotted. **D N-MNIST** [55] **sample;** original symbol digit: 7; 179 events. **E ebtMNIST-D1 sample;** original symbol’s digit:8; 35 events. **F ebtMNIST-D2 sample;** original symbol’s digit: 3; 16 events. **G Histogram of the time it took the EB camera to generate 48 events** starting from t=100 ms (100 ms after motor command was sent) for the 60K ebtMNIST-D1 dataset training set samples. **H Inter-Event Interval (IEI) histogram.** Histogram of time differences between any two consecutive events over all the pixels in the sensor. **I Cropped frames acquired with the event-base camera** of four MNIST symbols at distance D1 (fbtMNIST-D1 dataset); size of cropped area is 10×10 pixels.

Under stable illumination, events are generated by the camera only if there is relative motion between the camera and the scene. Similarly to [55] we opted for leaving the scene static and only moving the camera to create a relative motion, in order to mimic biological vision and avoid artifacts related to displaying motion on a digital display. The camera’s motion amplitude and velocity, as well as the size of the symbols on the sensor, were set to match relevant biological parameters (Supplementary Table S1).

### 2.2 Accurate Event Timing Contains Task-Relevant Information

To assess whether extra information is contained in precise timing of events we used a PointTransformer (PTr)[61] classifier with ∼ 5.9*M* to ∼ 11.7*M* parameters (see Methods for implementation details). The model receives a time series of events (analogous to retinal ganglion cells’ spikes) presented as tuples of (t, x, y, p), with t being time, x and y corresponding to the spatial coordinates on the photosensor array and p being polarity. This model enables access to temporal information with arbitrary precision and without any temporal information loss. PTr performs slightly better than discrete time recurrent architectures (GRU; ∼ 5.6*M* parameters; Supplementary Fig. S1 A). The FB data was evaluated using ResNet-18 (11.7M parameters).

The accuracy of the PTr classifier as a function of the number of input events saturated rapidly (Fig. 2A,B). For both D1 and D2 datasets, accuracy approached saturation within 70 to 100 events, with D2 datasets reaching saturation more quickly (Fig. 2A,B). During camera motion, the mean event generation rate was approximately 2.5 events/ms for ebtMNIST-D1 (Fig. 1B) and 4.0 events/ms for ebtFMNIST-D1, while for ebtMNIST-D2 and ebtFMNIST-D2, the rates were around 1.5 and 2.3 events/ms, respectively.

**Figure 2:**
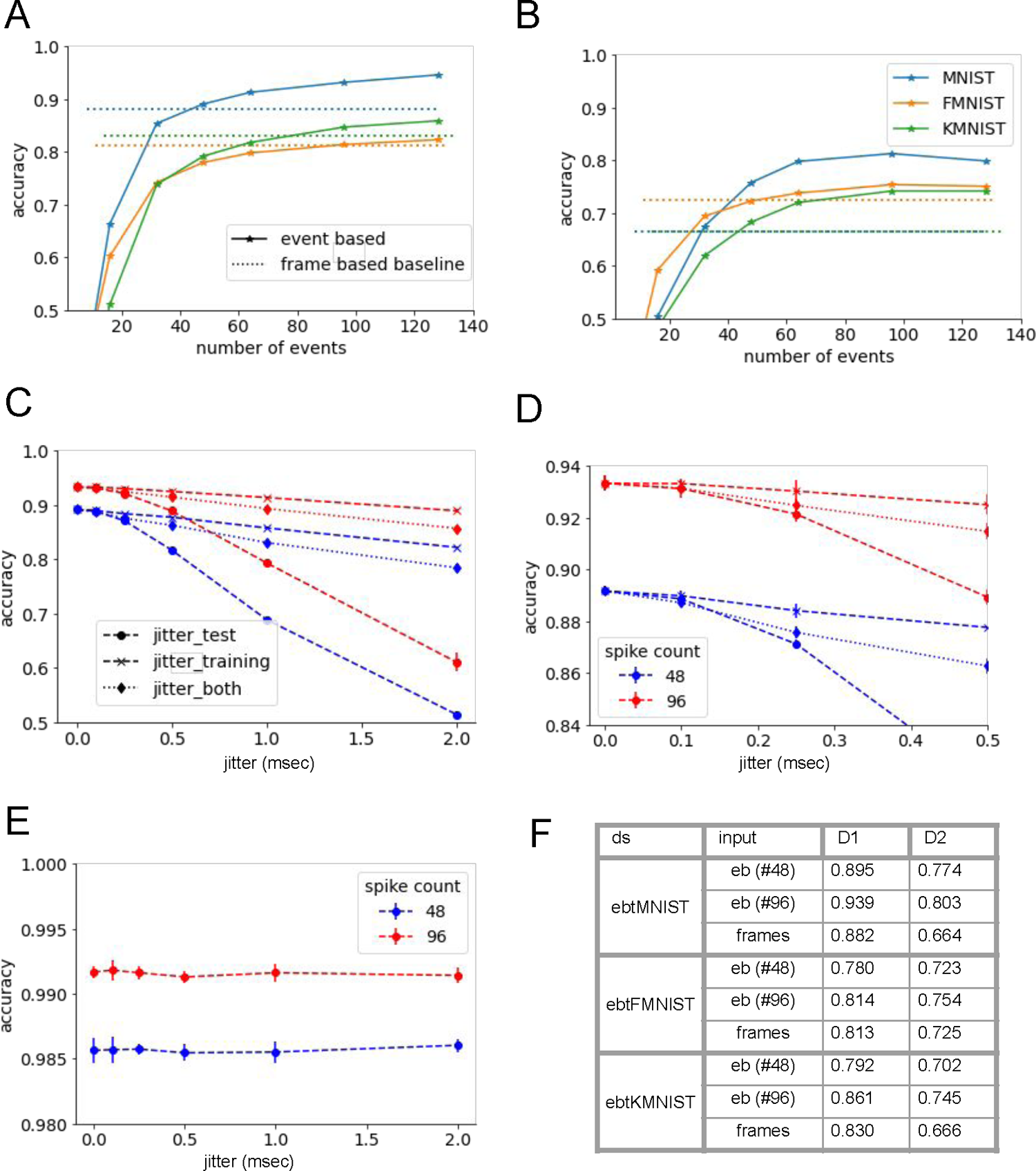
**A,B** Classification accuracy for tiny image datasets is shown vs number of events, for acquisition distance D1 (A) and D2 (B). The dashed lines denote FB classification accuracy (Resnet-18), for the corresponding datasets and conditions. **C,D Importance of temporal precision for tiny images**: MNIST at distance D1 is evaluated in presence of temporal jitter added to either at training, testing, or both stages of learning. **E Unimportance of temporal precision for large images**: lack of effect from jitter is demonstrated for N-MNIST dataset (see Fig. 1 for representative spatio-temporal raster plots of tiny and large stimuli). **F** Information from panels A,B presented in tabular form.

For almost all conditions the accuracy of our model on EB data surpassed that of a standard CNN trained on the equivalent FB datasets within less than 100 events. For example, with ebtMNIST-D1, our baseline model (PTr, 5.8M parameters) exceeded the accuracy of a standard CNN trained on fbtMNIST-D1 with just 48 events (Fig. 2A), achieving a test-set accuracy of 0.891 ± 0.002 compared to 0.882 ± 0.001 for the FB network (Fig. 2F). Based on this result, we selected 48 events as the default input size for all further experiments unless stated otherwise. Throughout the paper, we use a fixed number of events as the controlled parameter rather than a fixed time interval. However, for the basic setting (ebtMNIST-D1), we also report accuracy as a function of the time interval (Supplementary Fig. S1B).

Accuracy monotonically increased with additional events (Figure 2A). Larger transformer model (∼ 11.7*M* parameters, similarly to ResNet-18) reached higher accuracy (0.918±0.001 for the same event count), but required contrastive pre-training to converge (see Fig. 3C, **Event streams Are Inherently Augmented and Suitable for Self-supervised Learning** below and Methods).

**Figure 3:**
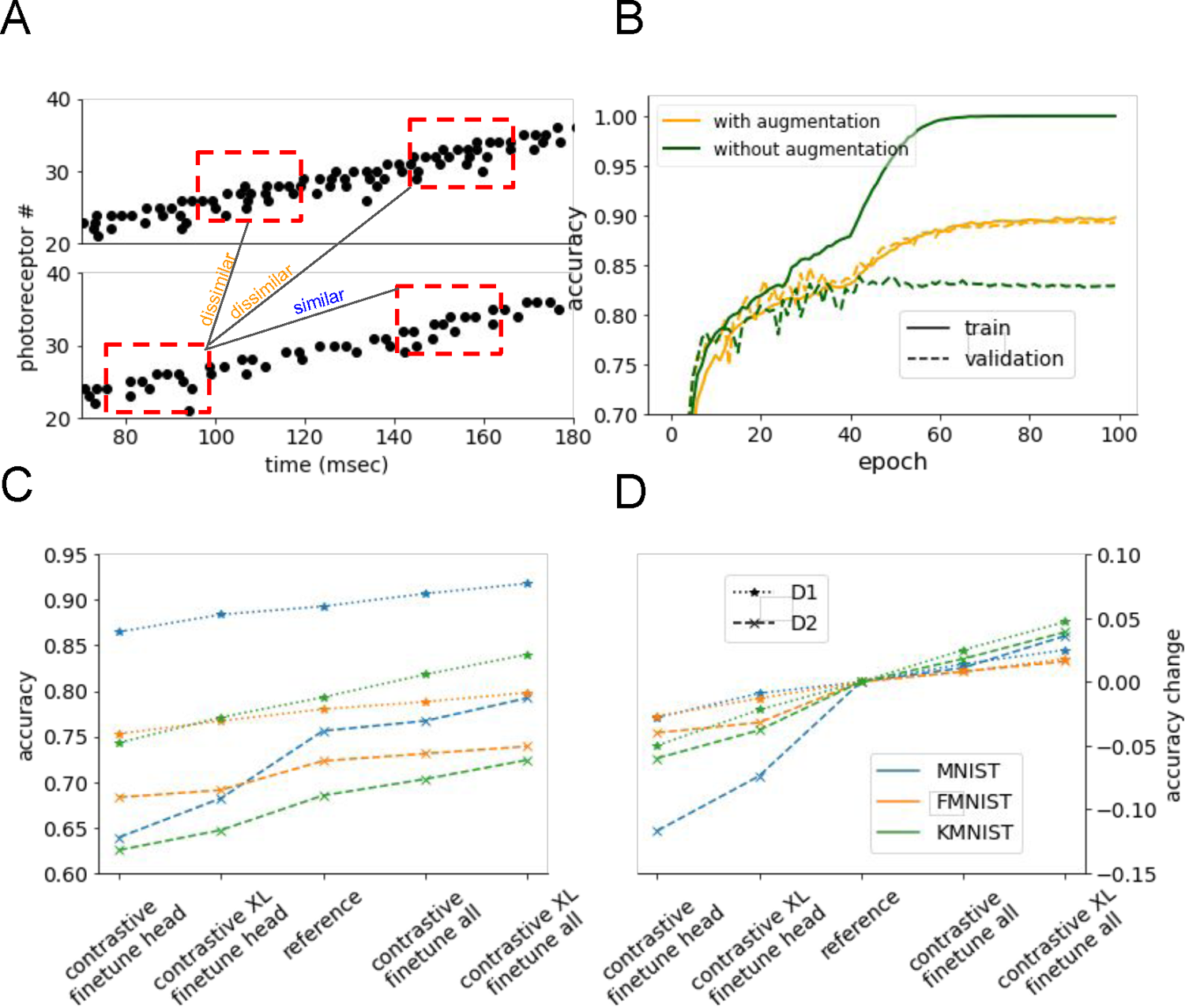
Data augmentation and contrastive learning with event sequences: **A** Contrastive learning is achieved by rewarding the similarity of neural representations across different time intervals of the same stimulus while discouraging similarity when the intervals correspond to different stimuli. Augmentation is performed by using variety of time intervals during training phase. Learning rate slowdown begins at epoch 40, which explains the abrupt rise in accuracy around this time. **B** Accuracy when learning from training data augmented by sequences from surrounding time-intervals compared to accuracy when restricting training data to the test interval. **C,D** Impact of contrastive pre-training is shown across datasets, network sizes and fine tuning methods. **C** absolute test accuracies. **D** differences to the reference classifier that is reported in Figure 1.

In order to asses the significance of the temporal accuracy of the sensor to classification accuracy we tested the performance of the transformer with additional artificial stochastic jitter added to the events. Added jitter values were sampled from a normal distribution with 0 mean and standard-deviation ranging from 0.1 ms to 2.0 ms (See Methods for more details).

Degradation in classification accuracy for the six datasets was clearly observed as temporal noise level was raised (Figure 2C,D, describes one representative dataset and supplementary figure Fig. S1 describes all datasets). This result indicates that the network has learned to extract the temporal information and utilize it for the classification task it was trained to solve. At jitter magnitude of 0.1 ms, which corresponds to ∼ 33% of the median inter-event interval (IEI, Fig. 1H; median is 0.33 ms), the impact of the added noise was negligible. However, increasing the noise to 0.25 ms (∼ 75% of the median IEI) led to significant degradation in classification accuracy (Figure 2C).

While degradation of accuracy for not-seen-before jitter during test is not surprising, the other two observations are more intriguing: First, we observed that applying jitter both to test and training lead to degraded accuracy, implying that the classifier failed to learn to overcome such noise. Next, we examined applying jitter only during training while testing the model with unperturbed samples. Here two opposing factors could theoretically play a role: jitter could augment the data by providing more training samples, and it could compromise the model’s ability to capture fine-grained timing details, which are essential for accurate classification according to active sensing models [14]. Our results support the latter effect: even a setting where jitter was added only to training data while test data was left intact led to degradation of classification accuracy. Thus, in our setting (in contrast to e.g. [62]), the overall impact of training with jitter was negative, namely degraded temporal information during training out-weighted the positive effect that the same jitter might have in augmenting the training set.

Conversely to additional jittered samples, higher event count, or, equivalently, longer acquisition time, did compensate for temporal jitter. For example, for ebtMNIST-D1, classification accuracy based on our default event number (48) with no added jitter (0.891±0.001; 2C) was approximately equivalent to processing 96 events with added temporal jitter with magnitude of 1.0 ms (0.894±0.002; 2C). It had been shown that human observers can mitigate spatial noise using longer viewing times [63, 64, 65, 66]. It would thus be intriguing to experimentally test whether temporal noise is similarly mitigated by longer fixations in humans, using settings with closed-loop gaze contingent displays [67, 68, 69].

We emphasize, however, that while our results support the claim that temporal information plays central role in classification of tiny images, it is redundant for full resolution images as was demonstrated by [53] for a wide range of datasets. We corroborated their finding for the pioneering event based dataset N-MNIST [55], which is based on full resolution MNIST, by repeating the experiment using N-MNIST samples. this experiment resulted with no apparent effect of added temporal jitter with the same magnitudes that affected the classification of tiny images (2E).

### 2.3 Event streams Are Inherently Augmented and Suitable for Self-supervised Learning

Unlike FB systems, which frequently process identical frames repeatedly, EB systems are continuously exposed to a dynamic, ever-changing stimulus, making each encounter with the stimulus truly “*Ichigo ichi-e*” — a unique, one-time experience. This inherent variability makes augmentation readily available and particularly effective. Here, by leveraging the temporal nature of event sequences, we augmented the training data by varying the time windows from which event sequences are extracted. Specifically, during training, we used event sequences with starting times shifted randomly, uniformly within the range of ±50 ms, from the original time window used for testing. This method led to approximately a 7% improvement in test accuracy compared to using a non-augmented dataset (Fig. 3A,B). As such augmentation capitalizes on the natural variability in EB data, it is expected to benefit both biological and computer vision agents. Both types of agents can use such augmentation to learn to perceive challenging scenarios, in which rapid perception is required and only a short glimpse of the scene is available.

To further harness the benefits of EB data, we developed an unsupervised pre-training method using contrastive learning (e.g., [70] and references therein). The key idea was to obtain task-relevant neural representations in an unsupervised manner, by enforcing consistency over time: bringing similar (positive) pairs closer together in the network’s neuronal space while pushing dissimilar (negative) pairs further apart. Positive pairs were generated by randomly selecting intervals from the same event sequence (we refer to these as time-based positive samples (TBPS)), while intervals from different stimuli served as negative pairs. This approach aligns with the rationale, which likely applies also to biological learning, that stimuli captured within short intervals (order of 200 ms) are affiliated with the same external object (Figure3A). We trained our model following standard contrastive learning procedures with a projection head (see Methods for details) in place of the classification head (e.g. [71]), with positive pairs obtained via TBPS. After this pre-training, we replaced the projection head with a classification head and conducted supervised training, training either the entire model or just the classification head, to optimize its performance as a classifier.

We evaluated the standard outcomes of contrastive training in our TBPS setting, focusing on three key aspects: (i) The amount of information contained in the resulting neural representation, determined by its ability to support a downstream task (classification, in our case) through the addition of a simple decoding head; this was quantified by comparing its performance to that of a fully supervised model. (ii) The extent to which TBPS-based contrastive pre-training improved performance when the entire model was fine-tuned. (iii) Whether TBPS-based contrastive pre-training enables the training of larger models that would otherwise fail to converge.

Fig. 3C,D presents the outcomes of TBPS-based contrastive pre-training across various datasets. The x-axis shows the baseline reference model (middle), with results displayed for TBPS-based contrastive pre-training followed by fine-tuning of the head alone (left) and full fine-tuning (right). In the head-only fine-tuning scenario, where only the classification head was trained while the rest of the model remained frozen, the results were only slightly inferior to those achieved with full fine-tuning (addressing question (i)), demonstrating the ability of the temporal contrastive pre-training to produce representations that are nearly sufficient for downstream classification tasks. TBPS-based contrastive pre-training further significantly improved performance when the entire model was fine-tuned (addressing question (ii)), confirming its utility in enhancing model capabilities. Furthermore, we evaluated an enlarged model, denoted as ‘XL,’ with a parameter count close to that of ResNet-18 (11.7M), previously used as a FB classifier in Fig. 2AB. For the XL model, TBPS-based contrastive pre-training proved essential, as the model failed to converge under vanilla supervised training (addressing question (iii)). These results highlight the dual advantages of TBPS-based contrastive pre-training in both constructing effective representations and enabling the training of larger architectures.

### 2.4 Testing for Vernier Hyperacuity

Humans can resolve Vernier offsets with hyperacuity [1, 2]. Whether brains resolve Vernier offsets using spatial or spatio-temporal information is debated [1, 21, 9]. We used our event based classifier for testing the feasibility of the spatio-temporal schemes.

In order to evaluate our classifier’s Vernier acuity we recorded events using the EB camera while it was presented with Vernier-like stimuli with offsets ranging from 1 to 7 screen pixels in both directions (Figure 4A,B,C). Following common psychophysical procedures, where subjects are first shown relatively large Vernier offsets and then tested on smaller ones, we checked how a Ridge linear regression performed on large Vernier offsets generalizes to smaller ones. That is, we used the Vernier stimuli with large offsets (6-7 pixels) to train the regression model and tested it on all Vernier stimuli (large and small offsets; 1-7 pixels). Similarly to the human Vernier task, the model task is to correctly report the direction of the offset, not the exact offset size. The accuracy of two regression models trained on top of the first and last transformer blocks were tested. The model trained on top of the last block exhibited hyperacuity for all tested offsets (the smallest of which was 1 pixel on the screen and thus ∼ 1*/*7 photosensor in the camera), resembling Vernier accuracy in humans (e.g. [72, 73]). Decoding from the first block yielded significantly lower performance (Figure 4D). These results indicate that fine, subpixel, visual details are enhanced rather than abstracted out as information passed from lower layers to higher ones.

**Figure 4:**
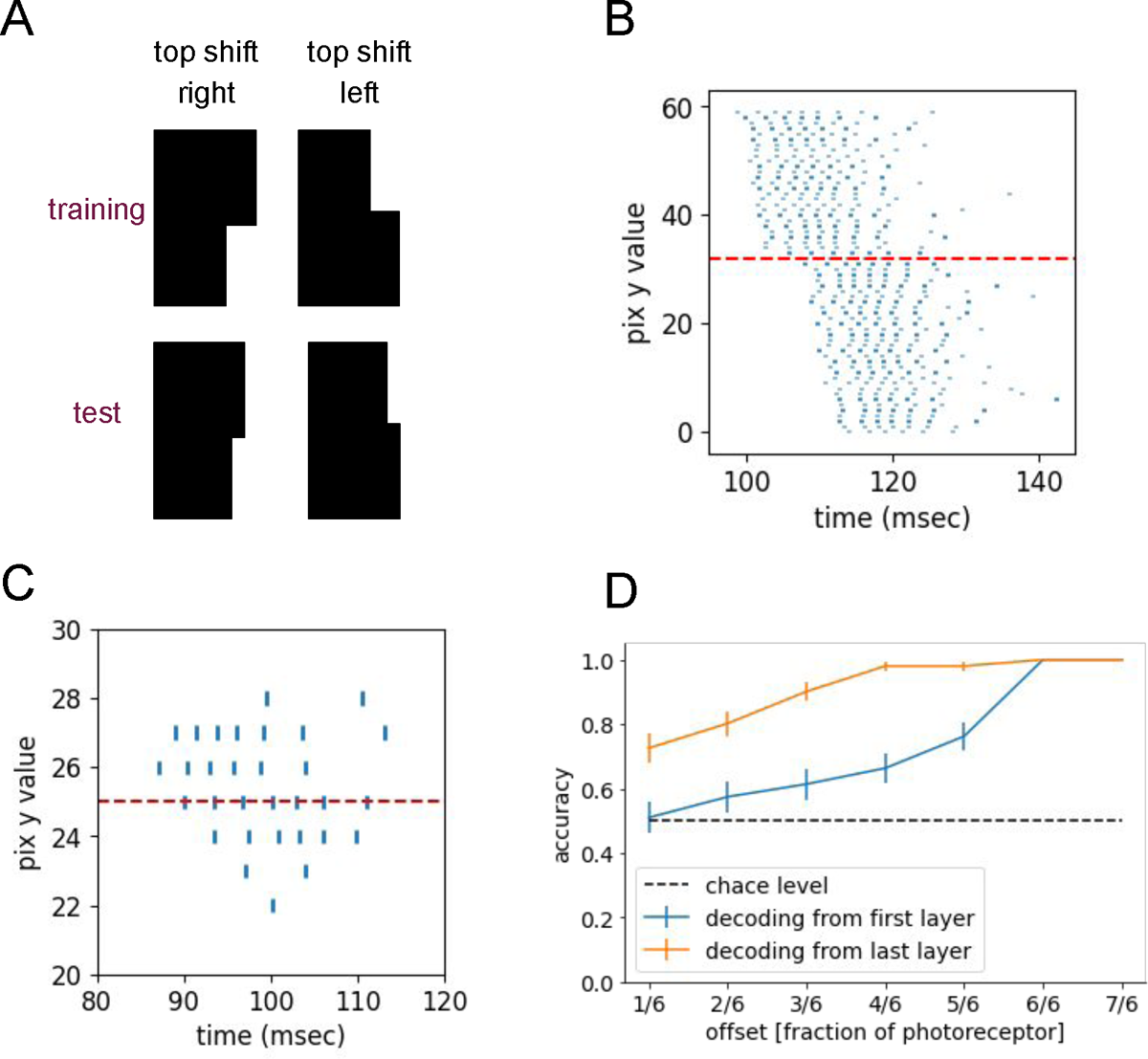
A Vernier task illustration: The agent is requested to report whether the top of the bar is shifted left or right with respect to its bottom. Training is done with large offsets and the testing is performed on small offsets. **B** Sensor activation raster plot shows events emitted by a 60 photosensors column when crossing a vernier stimuli like the one presented in (A; top-left) while rotating to the right. Numbers on the y axis correspond to photosensor’s index within the column. Dashed red line corresponds to the vertical position of the vernier bar shift. Events below and above the red dashed line are shifted by less than 10 ms, which corresponds to sensor speed of ∼ 30 deg/s. Events are not perfectly aligned in time across photosensors due to sensory and motor noise; single crossing of the edge results with generation of bursts of events which last ∼ 15 ms. **C** Same as (B) but for small edge where edge’s vertical dimension on display is equivalent to a single MNIST symbol (28 pix), which corresponds to ∼ 5 photosensors at D1 distance. Examples depicted at (B) and (C) correspond to an offset of a single photosensor (large offset) for the sake of clarity.**D** Accuracy in vernier task with small edges (see panel (C)). Reading out from first and last transformer block in an ANN trained as a classifier of ebtMNIST (at D1). Offsets of 6-7 pixels were used as training data and offsets of 5-1 pixels were used as test data. N=50 examples of each offset were presented.

## 3 Discussion

When challenged with tiny objects, classification that is based on purely spatial information is limited by the resolution of the photosensor array and is impaired by its motion (Fig. 1E,F). Here we showed that when spatiotemporal rather than spatial coding is used, sensor motion becomes a feature rather than a bug [9, 74]. As a result, Vernier hyperacuity emerges and recognition of tiny images is improved. Importantly, spatiotemporal processing is crucial in cases where the sensory information is limited, such as with tiny objects; when presented with large enough images, spatial processing is often sufficient [53].

Our results, obtained using a robotic bio-inspired vision system, are consistent with the claim that biological visual processing relies on both temporal and spatial information [75, 19, 8]. Specifically, we demonstrated that longer time spent looking at an image improves classification accuracy, which aligns well with human studies[63, 64, 65, 66]. Furthermore, we replicated a textbook experiment for probing hyperacuity in our setting, showing that training on tiny images is sufficient for resolving Vernier’s stimuli with human-like hyperacuity [73].

Precise temporal information obtained through EB vision was shown to contribute to the perception of dynamic scenarios such as action-recognition [53], to optical-flow computation [26, 76], and to 3D reconstruction from stereo-vision [77, 78]. Temporal precision was also shown to be beneficial for overcoming smearing by allowing motion compensation or motion cancellation [79, 80]. However, despite the demonstration that precise timestamps of visual events generated using static images contain information about the presented visual pattern [51], it has been commonly assumed that EB cameras are a poor choice for the perception of static scenes [54]. This has been based primarily on the fact that the EB signal is generated by relative motion; hence, with static scenes the camera has to be moving and this motion results with a distorted image of the static scene. Indeed, in the few cases where large EB static image datasets (N-MNIST, N-Caltech101 [55]) were used for image recognition [81, 82, 83, 30, 84, 31], spatial processing resulted with equivalent or superior performance compared to the proposed EB spatio-temporal models [53].

Our results disagree with the general assumption stated above and deviate from those showing that fine temporal resolution contributes mainly to precise motion compensation [54], for example for ‘canceling’ motion and generating static representations of a scene [13]. In our case, precise temporal information was used to build representations rather than to cancel their perturbations and by doing so we achieved superior performance compared with the equivalent spatial frame-based models.

Relevance of short timescale is further emphasized by our observation that the saturation of accuracy over event count in case of large images (N-MNIST) is rapid and follows the same pattern as for tiny images (supp Fig. S1C). This indicates that the relevant event count for performance is of the order of tens, which spans a time interval in the timescale of 1 ms.

The model we used, a PointTransformer [61], is not biomimetic. We used its ability to process sequences of spatio-temporal tuples, used here to process event streams while preserving accurate temporal information, for assessing the value of precise temporal information for image recognition. For further understanding of biological vision future work can extend the findings presented here to models with increased levels of biomimicry, including asynchronous networks (SNNs) and biomimetic learning rules, using neuromorphic processors [85, 86, 87].

With spatiotemporal coding, the resolution of encoding external space is determined by the combination of spatial and temporal factors. The spatial limiting factors are the hard-wired geometry of the photosensor array and its optics. The temporal limiting factors are the temporal precision of the photosensors and sensor motion speed (see Figure 4C for an illustration). When exploiting the temporal coding domain, biological agents can adjust sensor speed to balance increased sensor resolution (achieved at lower speeds) with reduced overall recognition time (achieved at higher speeds) [69, 88]. In our experiment with ebtMNIST-D1, our default event count of 48 events was typically acquired over time interval of around 20 ms (Figure 1G) with horizontal velocities of ∼30 deg/s (Figure 1C). Hence, symbols were typically shifted of by more than 2 pixels on the sensor during the acquisition of default event count, which corresponds to ∼ 70 − 80% of the tiny symbol size.

Humans exhibit ocular drifts at instantaneous speeds around 1 to 10 deg/s [8]. Assuming foveal receptive fields of the order of 0.01 deg in diameter [89], the temporal delays generated by a single edge crossed by neighboring receptors are in the order of 1 to 10 ms. Assuming a neuronal temporal resolution of 1 ms [15, 90], spatial offsets as small as 0.1 of the diameter of a single photoreceptor can be resolved. This is indeed demonstrated in our setting, when imitating principles of human vision (Fig. 4; Table S1) using sub-millisecond temporal resolution. The dependency on high temporal resolution is evident from our specific findings, which indicate that additional temporal jitter as small as 0.5 ms results in significant reduction in accuracy (Figure 2C,D).

Recent experiments indicate that the ocular drift is not fully stochastic, and its speed and curvature are adapted to the visual stimulus [69, 68]. Our data, collected in an open-loop, stimulus-independent manner, can help the design of closed-loop visual systems. In such closed-loop systems, camera’s motion is part of the perceiver-environment dynamics, in which motion optimizes sensory and perceptual efficiency. In our setting, motion and sensory parameter values overlap in part with those typically employed by human observers (Supplementary Table S2). The advantages and limitations of humanlike parameter regimes can be tested in future biomimetic applications of closed-loop vision.

Our work also shows how an event based representation enables unsupervised, contrastive, learning. This approach leverages the inherent variability of subsequences within an event sequence related to a static object (Fig. 3A) to facilitate learning (Fig. 3C,D). Specifically, our paradigm used different spike trains, generated during a single symbol acquisition, to train internal representations of the same category (Fig. 3). It is reasonable to expect that, similarly to our paradigm, the brain uses prior knowledge of the approximate constancy of the environment over tens of ms to develop useful representations. Future work could extend this unsupervised approach to train a generic EB model without requiring extensive labeled datasets.

### 3.1 Concluding remarks

We revealed two previously hidden advantages of EB sensing: its ability to perceive tiny images through precise temporal coding and its effectiveness in training. We demonstrated these advantages on relatively simple datasets, highlighting the need to extend this research to more complex and diverse real-world visual tasks. Furthermore, we analyzed the similarities between artificial EB sensing and biological vision, proposing that both rely on similar principles—a claim supported by the resemblance we demonstrate between human and machine hyperacuity.

## 4 Methods

### 4.1 Setup

An EB camera (DAVIS240C, iniVation AG; [57]) was used to generate event streams of small static objects while rotating. In order to allow controlled rotations the camera was mounted on a pan-tilt unit (PTU). The PTU was composed of 2 servo motors (TGY-D003HV, TURNIGY), a small motion controller (Mini Maestro, Pololu) and mechanical parts that were produced in-house. A computer display was positioned in front of the camera 1A. This setting was inspired by [55].

The camera, the controller and the display were connected to a single computer that was used to save the camera’s output (events/frames, Inertial Measurement Unit (IMU)), initiate and end camera’s cyclic rotation motion, and display the symbols. Camera’s IMU was used to continuously record the angular velocity of the camera at a sampling rate of 1KHz (Figure 1C). One of the controller’s digital output was connected to the trigger-in input of the camera in order to accurately synchronize the beginning and end of rotation movements with the visual events generated by the camera.

Code written in python was used to coordinate the display of the symbols, the motion of the camera and the acquisition of files containing the camera’s output. We used the dv-runtime software (iniVation AG) to initialize the camera, set its parameters and retrieve its output. Data generated by the camera was saved using this framework to files on the computer.

### 4.2 Datasets acquisition

Our setup was used to convert popular image classification datasets to EB and FB tiny images datasets. Source datasets’ symbols were projected on the computer display at their original resolution of 28×28 pixels (Fig. 1). Datasets were recorded with the camera positioned at one out of two possible distances from the display: 525 mm (configuration ‘D1’) and 830 mm (configuration ‘D2’). Configuration ‘D1’ resulted with symbol covering ∼5×5 sensor pixels while at configuration ‘D2’ symbol projected size was ∼4×4 pixels (see further details regrading projection size below).

The datasets were acquired using simple motion profiles, involving either a single fixed-velocity horizontal rotation (pan) or a fixed-velocity pan followed by a fixed-velocity vertical rotation (tilt). Notably, while the nominal speed was fixed, the actual speed varied significantly as can be seen in Fig. 1C.

Illumination conditions were kept fixed during all dataset acquisition sessions. The setup was placed within a carton box in a room lighted by led lighting and with window curtains closed. This resulted with environment with steady dim illumination during all hours as the main source of illumination reaching the camera was the computer display. Display background was set to 20% brightness while stimuli pixels values (0-255) were linearly mapped to the 20%-100% grey level range.

Camera’s biases were kept fixed as well for all acquisition sessions (dv-runtime configuration file that includes camera biases is included in the Github repository). For FB datasets, camera’s frame-rate was set to 20 Hz while exposure time was 4 ms.

Data acquired was first saved by the dv-runtime in aedat4 format. Pre-processing of the data, before it was used to train the networks, included parsing the long stream of events to a list of streams where each such stream contains events that where generated during the presentation of a single symbol, and saving the parsed data to a file using a different binary format using the Python pickle library.

The order of acquisition was set randomly and shuffling was re-applied for each acquisition session. The test samples (10K) were embedded randomly within the sequence of train samples (60K) during acquisition, in order to mitigate the potential effect of biases developing along the period of single dataset acquisition.

### 4.3 Acquisition of EB Vernier data

Acquisition of several EB datasets incorporated also Vernier stimuli. In this cases Vernier stimulus was presented on the computer display once every 100 repetitions. Camera motion and configuration during the Vernier repetition was identical to the ones during acquisition of the source dataset symbols (MNIST, FMNIST or KMNIST). This guaranteed acquisition with same conditions for Vernier as for the other tiny-symbols datasets. Vernier edge vertical dimension was set to 28 display pixels to make it spatially comparable to the other symbols displayed and sampled by the EB sensor. For each Vernier repetition the horizontal offset in display pixels was chosen from the range of ±1 to ±7 with plus or minus sign, where the sign determined offset direction. In total the complete Vernier dataset included at least 50 repetitions for each of the offsets that were used.

### 4.4 Estimating the display to photosensor-array projection ratio in pixels

We computed the size of a single display pixel projected on the sensor in sensor pixels for the two distances used to acquire the datasets (D1 and D2). In order to do so we used the camera’s magnification formula:

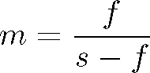

where *m* is the magnification, *f* is the focal length of the lens and *s*is the distance between the computer display and the camera. In order to get the ratio in units of *pix_display_/pix_sensor_* we multiplied ( m) by the *pitch_display_//pitch_sensor_*ratio where *pitch* is the size of a single pixel. Given that a 4.5 mm focal-length lens was used, the DAVIS240C camera pixel size is 18.5 *µ*m and the used display pitch was 0.275 mm, the display pixel size to sensor pixel size ratio was 0.129 and 0.081 for D1 (s=525 mm) and D2 (s=830 mm) respectively. We further estimated the ratio for D1 empirically. This was done by measuring the distance in sensor pixels between two vertical lines displayed at a known horizontal distance on the display and projected onto the sensor, while camera is positioned at the D1 distance. Using this method we found that a single display pixel was mapped to ∼ 1*/*8 sensor pixel for the D1 configuration (data not shown). The estimated ratio of ∼ 1*/*8 is consistent with the calculation.

These ratios are not consistent, however, with the numbers reported above as the ∼20×20 pixels of the original MNIST symbols on the display were projected to 5×5 pixels on the sensor. the extended projection size on the sensor ∼ 5 pixels was caused by the non-ideal lens aberrations which resulted with de-focusing of the ∼ 2.5 pixels expected ideal sharp image.

### 4.5 Model implementation and training details

#### Transformer-based Model

Transformer-based architecture, similar to Point Cloud Transformer [61], was employed for time-series classification. Input dimensionality was four, rather than three features per event (’point’) to account for two spatial dimensions (event’s x,y position on the sensor), one temporal dimension (event’s timestamp) and one polarity bit (event’s polarity). First, each feature was shifted and scaled element-wise by fixed offsets (−110, −35, −26, 0) and scalings (0.1, 1.0, 1.0, 1.0), this resulted with approximately normalized data. We chose to use this fixed approximate normalization rather than per batch or per sample normalization in order to preserve relative spatio-temporal information. These processed inputs were then projected from dimension 4 to 256 via a learnable linear transformation.

The projected features were fed to *l* = 8 successive Transformer layers, each composed of a multi-head self-attention module with 4 heads (key/query dimension *d_k_* = 256) and a position-wise MLP of dimension 512. Residual connections and layer normalization were applied around both the attention and feedforward modules. A dropout rate of 0.1 was used throughout the self-attention and feedforward sub-layers, starting from layer 4.

For the final classification, we adopted an average-pooling head that computed the mean across events and then applied a fully connected projection of dimension 256. After another dropout step, a final linear layer produced logits for 10 output classes.

#### GRU-based model

We used plain multilayer GRU with 14 layers to approximately match the parameter count of the PTr version. All other design choices were identical to the baseline PTr model: Input constituted event’s x,y position on the sensor, timestamp and polarity. Classification head was identical to the one used with the PTr version and relied on average activation of the last GRU layer over all time-steps (i.e. average pooling). Input pre-processing, which included centering and scaling, was identical also to the pre-processing used with the baseline model.

#### Contrastive learning

For contrastive pretraining [71] the classification head was replaced with projection head that ended with projection into a 256 dimensional embedding space rather than into 10 classification space. It is common practice [71] that a projection head is a temporary set of neural network layers that transforms the features learned by the model into a space where similar examples are pulled together and dissimilar ones pushed apart—helping the model to learn meaningful patterns without relying on labels. After this pre-training phase, we replaced the projection head with a classification head, which is the final set of layers designed to interpret these learned features and assign each input to a specific category. We then conducted supervised training, updating either the entire model or just the classification head, to optimize its performance as a classifier. Normalized Temperature-scaled Cross Entropy Loss with temperature of 0.2 was used to train the model. In this method cross entropy loss is applied to logits that are computed as cosine similarities (scaled by temperature) between samples (see [71] for details). We observed empirically that for optimal performance the ratio between positive and negative pairs should be small relative to ratios reported in literature [71]. Positive/negative ratio was set to 5/64 in average, which was achieved using batches of 64 with 4/5 of negative samples randomly dropped (such dropping was more time-efficient than working with smaller batches).

#### Optimization schedule

The model was trained for 100 epochs, using a batch size of 64. We used Adam optimizer with a max learning rate of 1 × 10*^−^*^4^, linear warmup of 5 epochs with initial rate of 1 × 10*^−^*^6^, and exponential decay of learning rate with factor 0.9 starting from epoch 40.

### 4.6 Adding temporal noise (jitter) to event-based datasets

We have added temporal noise to acquired data in order to estimate its effect on network’s performance. The added noise for a single training instance was sampled from a normal distribution with 0 mean and standard-deviation chosen within the range of 0.1 to 10.0 ms.

When applied to training data jitter values were sampled once for each event and used along the training (i.e. jitter was not resampled for each training epoch the way it is usually done when applying data augmentation, since we wanted to simulate higher noise level at the sensor).

### 4.7 Experimental repetitions, data splits and Outlier Exclusion Criteria

Five random seeds (i.e., variations in network initialization parameters and train/validation splits) were used to generate 5 training instances in all experiments. Development was preformed with validation set constituting 10% of training samples. Trained models for which test accuracy were reported were trained excluding validation set (extra 10% that might have been used for training in these cases) to ensure consistency with the development results. The classification accuracy numbers mentioned in the paper correspond to the Mean±Standard-deviation (STD) of the test set accuracy over these 5 different trained model instances of each experiment.

We excluded from our plots and results any seeds whose performance was more than 10% lower than the best seed’s accuracy or for which optimization collapsed; this 10% relative threshold corresponds to an absolute accuracy drop of at least 5%. In all remaining cases, the standard deviations across the reported repetitions was below 0.5%, clearly identifying the discarded instances as outliers. Across the 35 settings shown in Table S2, seeds were discarded in only four instances. Furthermore, among 30 settings in Table S4, there was only one case where a single seed was excluded.

### 4.8 Data availability

The datasets acquired and used during this study (including tiny event-based and tiny frame-based versions) are publicly available at datasets-link and can be used to train the models described above. Any other data will be shared by the authors upon request.

### 4.9 Code availability

The code used during this study is publicly available at https://github.com/ahissar-lab/event-based-hyperacuity. The code can be used freely for academic research.

## Acknowledgments

We thank Jonathan Victor and David Burr for commenting on earlier versions of the manuscript. This research was supported by the European Research Council (ERC) under the EU Horizon 2020 Research and Innovation Programme (grant agreement No 786949), the MBZUAI-WIS Joint Program for AI Research, the Weizmann institute sustainability and energy research initiative and the USA Air Force Office of Scientific Research (AFOSR, grant No. FA9550-22-1-0346, EA).

## Author contributions

El.A. and A.R. conceived the study, designed the study, collected the data, conducted the experiments, analyzed the data and wrote the paper. M.K. built the setup and collected the data. F.S.K. and S.K. advised on data analysis and wrote the paper. Eh.A. designed and supervised the study and wrote the paper.

## 5 Supplementary

**Table S1:** Comparison of metrics between Human Eye and DVS Sensor, the values are taken for sequence of 48 events at camera distance D1.

| Metric | Human Eye | DVS Sensor | Comments |
| --- | --- | --- | --- |
| Sensor/receptor size [arcmin] | 0.5 [89] | 15 | For camera we used:<br>$\frac{60 \text{ deg field of view}}{240 \text{ sensors}} = 15'$ |
| Acquisition time range [ms] | 25 - 100 [65] | 20 - 80 | Fig. 1G for DVS data |
| Typical velocity [arcmin/sec] | 55 [91] | 1800 | Fig. S9 in [91] for humans.<br>Fig. 1C for DVS data |
| Typical velocity [receptors/sec] | 110 | 120 |  |
| Typical number of crossed receptors during acquisition [receptors] | 2.25-11.10 | 2.4 - 9.6 | $\frac{1800' \text{sec} \times 20 \text{ms}}{15'}$ |

**Table S2:** Accuracy over event count: tabular results.

| Dataset | Distance | N<br>events | N<br>Runs | Accuracy<br>Mean | Accuracy<br>STD | Accuracy<br>SEM |
| --- | --- | --- | --- | --- | --- | --- |
| MNIST | D1 | 8 | 4 | 0.396 | 0.002 | 0.001 |
| KMNIST | D1 | 8 | 5 | 0.333 | 0.001 | 0.001 |
| FMNIST | D2 | 8 | 5 | 0.404 | 0.003 | 0.002 |
| KMNIST | D2 | 8 | 5 | 0.301 | 0.004 | 0.002 |
| FMNIST | D1 | 8 | 5 | 0.405 | 0.003 | 0.001 |
| MNIST | D2 | 8 | 1 | 0.305 | - | - |
| MNIST | D1 | 16 | 5 | 0.663 | 0.003 | 0.001 |
| KMNIST | D1 | 16 | 5 | 0.510 | 0.001 | 0.000 |
| FMNIST | D2 | 16 | 5 | 0.592 | 0.002 | 0.001 |
| FMNIST | D1 | 16 | 4 | 0.602 | 0.002 | 0.001 |
| KMNIST | D2 | 16 | 5 | 0.483 | 0.002 | 0.001 |
| MNIST | D2 | 16 | 5 | 0.504 | 0.004 | 0.002 |
| FMNIST | D1 | 32 | 4 | 0.742 | 0.001 | 0.000 |
| KMNIST | D2 | 32 | 5 | 0.619 | 0.002 | 0.001 |
| KMNIST | D1 | 32 | 5 | 0.738 | 0.001 | 0.000 |
| MNIST | D1 | 32 | 5 | 0.854 | 0.001 | 0.001 |
| MNIST | D2 | 32 | 5 | 0.675 | 0.004 | 0.002 |
| FMNIST | D2 | 32 | 5 | 0.694 | 0.002 | 0.001 |
| MNIST | D2 | 48 | 5 | 0.758 | 0.005 | 0.002 |
| KMNIST | D1 | 48 | 5 | 0.792 | 0.004 | 0.002 |
| FMNIST | D1 | 48 | 5 | 0.780 | 0.002 | 0.001 |
| KMNIST | D2 | 48 | 5 | 0.683 | 0.003 | 0.001 |
| MNIST | D1 | 48 | 5 | 0.891 | 0.002 | 0.001 |
| FMNIST | D2 | 48 | 5 | 0.723 | 0.002 | 0.001 |
| KMNIST | D1 | 64 | 5 | 0.818 | 0.002 | 0.001 |
| MNIST | D1 | 64 | 5 | 0.913 | 0.002 | 0.001 |
| KMNIST | D2 | 64 | 5 | 0.720 | 0.004 | 0.002 |
| FMNIST | D2 | 64 | 5 | 0.738 | 0.003 | 0.001 |
| FMNIST | D1 | 64 | 5 | 0.798 | 0.001 | 0.000 |
| MNIST | D2 | 64 | 5 | 0.798 | 0.004 | 0.002 |
| MNIST | D1 | 96 | 5 | 0.932 | 0.003 | 0.002 |
| FMNIST | D1 | 96 | 5 | 0.814 | 0.001 | 0.001 |
| KMNIST | D2 | 96 | 5 | 0.742 | 0.006 | 0.003 |
| KMNIST | D1 | 96 | 5 | 0.847 | 0.001 | 0.001 |
| FMNIST | D2 | 96 | 5 | 0.754 | 0.003 | 0.001 |
| MNIST | D2 | 96 | 5 | 0.813 | 0.004 | 0.002 |
| FMNIST | D1 | 128 | 5 | 0.823 | 0.002 | 0.001 |
| FMNIST | D2 | 128 | 5 | 0.751 | 0.004 | 0.002 |
| MNIST | D2 | 128 | 5 | 0.799 | 0.004 | 0.002 |
| MNIST | D1 | 128 | 5 | 0.946 | 0.002 | 0.001 |
| KMNIST | D1 | 128 | 5 | 0.859 | 0.002 | 0.001 |
| KMNIST | D2 | 128 | 5 | 0.742 | 0.002 | 0.001 |

**Table S3:** Adding temporal jitter: tabular results.

| mode | N<br>events | Jitter [ms] | N<br>Runs | Mean | STD | SEM |
| --- | --- | --- | --- | --- | --- | --- |
| jitter test | 48 | 0.000 | 5 | 0.892 | 0.002 | 0.001 |
| jitter test | 48 | 0.100 | 5 | 0.889 | 0.002 | 0.001 |
| jitter test | 48 | 0.250 | 5 | 0.871 | 0.002 | 0.001 |
| jitter test | 48 | 0.500 | 5 | 0.817 | 0.004 | 0.002 |
| jitter test | 48 | 1.000 | 5 | 0.689 | 0.005 | 0.002 |
| jitter test | 48 | 2.000 | 5 | 0.514 | 0.005 | 0.002 |
| jitter test | 96 | 0.000 | 5 | 0.933 | 0.003 | 0.001 |
| jitter test | 96 | 0.100 | 5 | 0.931 | 0.003 | 0.002 |
| jitter test | 96 | 0.250 | 5 | 0.921 | 0.003 | 0.001 |
| jitter test | 96 | 0.500 | 5 | 0.889 | 0.003 | 0.001 |
| jitter test | 96 | 1.000 | 5 | 0.793 | 0.007 | 0.003 |
| jitter test | 96 | 2.000 | 5 | 0.610 | 0.017 | 0.008 |
| jitter training | 48 | 0.000 | 5 | 0.892 | 0.002 | 0.001 |
| jitter training | 48 | 0.100 | 5 | 0.890 | 0.002 | 0.001 |
| jitter training | 48 | 0.250 | 5 | 0.884 | 0.003 | 0.001 |
| jitter training | 48 | 0.500 | 5 | 0.878 | 0.001 | 0.000 |
| jitter training | 48 | 1.000 | 5 | 0.858 | 0.002 | 0.001 |
| jitter training | 48 | 2.000 | 5 | 0.822 | 0.001 | 0.001 |
| jitter training | 96 | 0.000 | 5 | 0.933 | 0.003 | 0.001 |
| jitter training | 96 | 0.100 | 5 | 0.933 | 0.002 | 0.001 |
| jitter training | 96 | 0.250 | 5 | 0.930 | 0.004 | 0.002 |
| jitter training | 96 | 0.500 | 5 | 0.925 | 0.004 | 0.002 |
| jitter training | 96 | 1.000 | 5 | 0.914 | 0.003 | 0.001 |
| jitter training | 96 | 2.000 | 5 | 0.889 | 0.003 | 0.001 |
| jitter both | 48 | 0.000 | 5 | 0.892 | 0.002 | 0.001 |
| jitter both | 48 | 0.100 | 5 | 0.887 | 0.001 | 0.001 |
| jitter both | 48 | 0.250 | 5 | 0.876 | 0.002 | 0.001 |
| jitter both | 48 | 0.500 | 5 | 0.863 | 0.003 | 0.001 |
| jitter both | 48 | 1.000 | 5 | 0.831 | 0.003 | 0.001 |
| jitter both | 48 | 2.000 | 5 | 0.785 | 0.001 | 0.000 |
| jitter both | 96 | 0.000 | 5 | 0.933 | 0.003 | 0.001 |
| jitter both | 96 | 0.100 | 5 | 0.931 | 0.001 | 0.001 |
| jitter both | 96 | 0.250 | 5 | 0.925 | 0.002 | 0.001 |
| jitter both | 96 | 0.500 | 5 | 0.915 | 0.003 | 0.001 |
| jitter both | 96 | 1.000 | 5 | 0.894 | 0.004 | 0.002 |
| jitter both | 96 | 2.000 | 5 | 0.857 | 0.003 | 0.001 |

**Table S4:** Contrastive learning: tabular results.

| Dataset | Distance | Mode | N<br>Runs | Accuracy<br>Mean | Accuracy<br>STD | Accuracy<br>SEM | Accuracy<br>Change |
| --- | --- | --- | --- | --- | --- | --- | --- |
| MNIST | D1 | contrastive finetune head | 5 | 0.865 | 0.002 | 0.001 | -0.028 |
| MNIST | D1 | contrastive XL finetune head | 5 | 0.884 | 0.002 | 0.001 | -0.009 |
| MNIST | D1 | reference | 4 | 0.893 | 0.002 | 0.001 | 0.000 |
| MNIST | D1 | contrastive finetune all | 5 | 0.907 | 0.001 | 0.001 | 0.014 |
| MNIST | D1 | contrastive XL finetune all | 5 | 0.918 | 0.002 | 0.001 | 0.025 |
| MNIST | D2 | contrastive finetune head | 5 | 0.639 | 0.007 | 0.003 | -0.117 |
| MNIST | D2 | contrastive XL finetune head | 5 | 0.682 | 0.004 | 0.002 | -0.074 |
| MNIST | D2 | reference | 5 | 0.756 | 0.006 | 0.003 | 0.000 |
| MNIST | D2 | contrastive finetune all | 5 | 0.767 | 0.002 | 0.001 | 0.011 |
| MNIST | D2 | contrastive XL finetune all | 5 | 0.792 | 0.002 | 0.001 | 0.036 |
| FMNIST | D1 | contrastive finetune head | 5 | 0.753 | 0.002 | 0.001 | -0.027 |
| FMNIST | D1 | contrastive XL finetune head | 5 | 0.767 | 0.002 | 0.001 | -0.013 |
| FMNIST | D1 | reference | 5 | 0.780 | 0.002 | 0.001 | 0.000 |
| FMNIST | D1 | contrastive finetune all | 5 | 0.788 | 0.001 | 0.001 | 0.008 |
| FMNIST | D1 | contrastive XL finetune all | 5 | 0.798 | 0.001 | 0.001 | 0.018 |
| FMNIST | D2 | contrastive finetune head | 5 | 0.683 | 0.002 | 0.001 | -0.040 |
| FMNIST | D2 | contrastive XL finetune head | 5 | 0.691 | 0.003 | 0.001 | -0.032 |
| FMNIST | D2 | reference | 5 | 0.723 | 0.002 | 0.001 | 0.000 |
| FMNIST | D2 | contrastive finetune all | 5 | 0.731 | 0.003 | 0.001 | 0.008 |
| FMNIST | D2 | contrastive XL finetune all | 5 | 0.739 | 0.003 | 0.002 | 0.016 |
| KMNIST | D1 | contrastive finetune head | 5 | 0.743 | 0.005 | 0.003 | -0.050 |
| KMNIST | D1 | contrastive XL finetune head | 5 | 0.771 | 0.002 | 0.001 | -0.022 |
| KMNIST | D1 | reference | 5 | 0.793 | 0.004 | 0.002 | 0.000 |
| KMNIST | D1 | contrastive finetune all | 5 | 0.818 | 0.003 | 0.001 | 0.025 |
| KMNIST | D1 | contrastive XL finetune all | 5 | 0.840 | 0.002 | 0.001 | 0.047 |
| KMNIST | D2 | contrastive finetune head | 5 | 0.625 | 0.003 | 0.001 | -0.060 |
| KMNIST | D2 | contrastive XL finetune head | 5 | 0.647 | 0.002 | 0.001 | -0.038 |
| KMNIST | D2 | reference | 5 | 0.685 | 0.001 | 0.000 | 0.000 |
| KMNIST | D2 | contrastive finetune all | 5 | 0.703 | 0.004 | 0.002 | 0.018 |
| KMNIST | D2 | contrastive XL finetune all | 5 | 0.724 | 0.003 | 0.001 | 0.039 |

---

**Figure S1:**
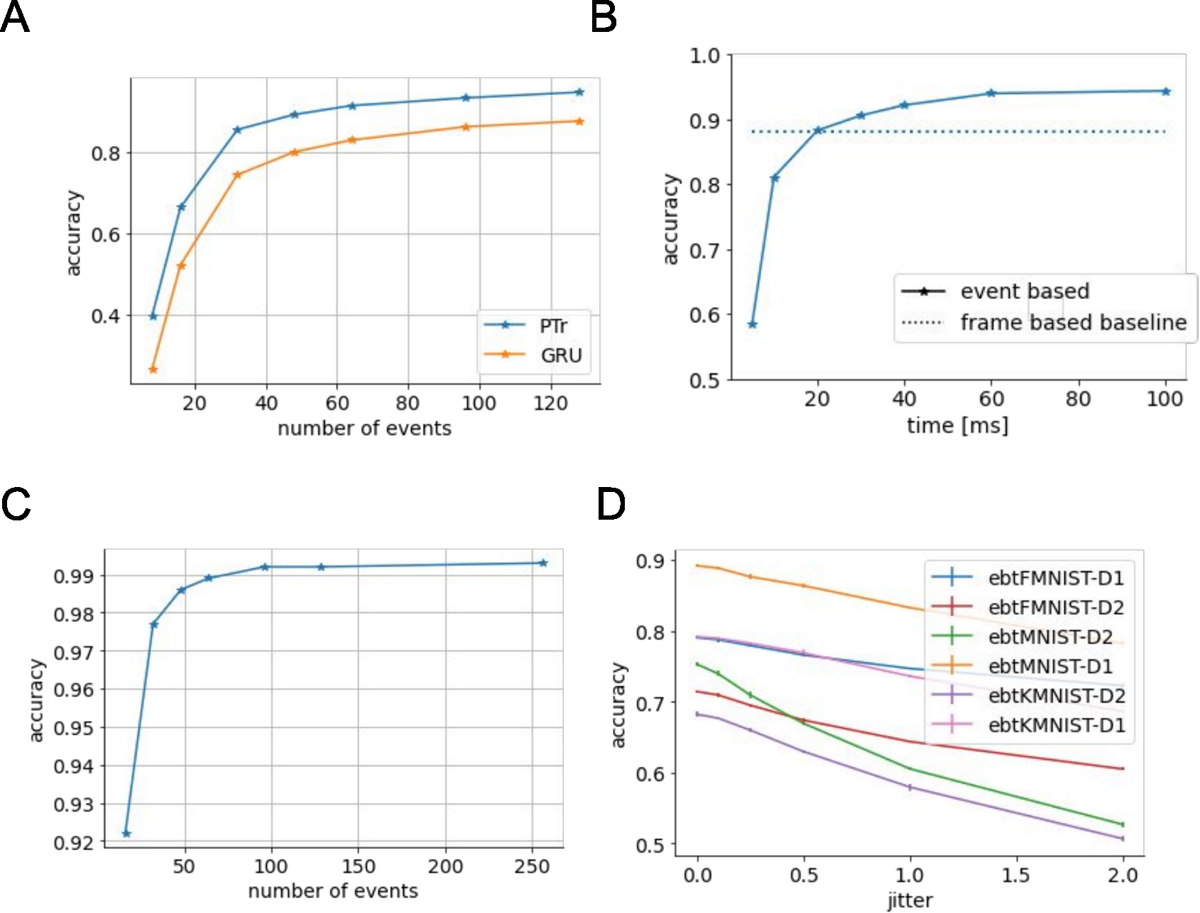
A. Point Transformer vs. Recurrent Neural Network. GRU (Methods for details) and Point transformer (PTr) with comparable number of parameters and training times are shown (parameter count: ∼ 5.9M for PTr, ∼ 5.6M for GRU). **B. Event based with controlled time window** Same as Fig. 2A, but with controlled parameter being events acquisition duration (i.e. all events occurred within a fixed time-window were use as input to the model) rather than event count. **C. N-MNIST** accuracy over different event counts. **D. Effect of temporal jitter** is shown here for all our datasets. The jitter was applied to both test and training sets

